# Estimating Individual Contributions to Complex DNA SNP Mixtures

**DOI:** 10.1101/391086

**Authors:** Darrell O. Ricke, Philip Fremont-Smith, James Watkins, Tara Boettcher, Eric Schwoebel

**Affiliations:** Bioengineering Systems & Technologies, Massachusetts Institute of Technology Lincoln Laboratory Lexington, MA USA

**Keywords:** Forensic science, DNA forensics, mixture analysis, single nucleotide polymorphism, high throughput sequencing, massively parallel sequencing

## Abstract

Mixture analysis and deconvolution methods can identify both known and unknown individuals contributing to DNA mixtures. These methods may not identify all DNA contributors with the remaining fraction of the mixture being contributed by one or more unknown individuals. The proportion of DNA contributed by individuals to a forensic sample can be estimated using their quantified mixture alleles. For short tandem repeats (STRs), methods to estimate individual contribution concentrations compare capillary electrophoresis peak heights and or peak areas within a mixture. For single nucleotide polymorphisms (SNPs), the major:minor allele ratios or counts, unique to each contributor, can be compared to estimate contributor proportion within the mixture. This article introduces three approaches (mean, median, and slope methods) for estimating individual DNA contributions to forensic mixtures for high throughput sequencing (HTS)/massively parallel sequencing (MPS) SNP panels.

## Introduction

The amount of DNA contributed by individuals to forensic DNA mixture samples is known to vary by individual(1, 2). Estimates of how much DNA an individual contributed to a mixture can provide useful insights into relative contributions by different contributors(1). Traditional STR forensics analysis leverages peak height or peak area information to estimate individual contribution amounts, deconvolute mixtures, and enhance likelihood ratio calculations(3–7). Recent advances are pushing the upper boundary to characterize STR mixtures with more contributors(8–15).

An alternative or complimentary approach for analyzing DNA mixtures is the characterization of SNPs using HTS/MPS technology(16). SNP panels have been designed to identify individuals in DNA mixtures with more contributors(17). Forensic SNP panels enable mixture deconvolution capabilities(18), expanded kinship identification(19, 20) (due to the larger number of loci characterized), biogeographic ancestry prediction(21, 22), and phenotype predictions(23). GrigoraSNPs(24), ForenSeq(25), ExactID(26), and STRaitRazor(27) are programs capable of characterizing MPS forensic SNP panels.

SNP panels have the potential to make significant impacts on forensic investigations. Similar to STR allele quantification, the ratios of major:minor alleles reflect DNA contributor concentrations. We introduce three methods (mean, median, and slope) for estimating contributor concentrations in mixture samples characterized by SNP panels(28) based on quantified major:minor alleles. These methods can quantify DNA contributions from multiple contributors, as well as estimate the proportion of DNA contributed by unknown contributors.

## Methods

### Ion Torrent MPS Sample Prep and Sequencing

For defined mixture experiments, swabs (Bode cat# P13D04) were used to collect buccal cells from the inside of cheeks of volunteers, rubbing up and down for at least 10 seconds, with pressure similar to that used while brushing teeth. For touch mixture experiments, swabs (Bode cat# P13D04) were used to collect DNA from objects of different surface types that had touch history logs. DNA was isolated from swabs using the QIAamp DNA Investigator Kit (QIAGEN cat#56504), using the “Isolation of Total DNA from Surface and Buccal Swabs” protocol, and eluted in 100uL of low TE (carrier RNA not used; low TE has 0.1mM of EDTA). Quantitation was done using Quantifilier HP kit (Thermo Fisher Scientific cat#4482911) according to manufacturer recommendations, with the exception that human genomic DNA from Aviva Systems Biology (cat#AVAHG0001) was used for the standard. Purified DNA from individuals were combined to produce defined two to ten person mixtures. Primers for the 2,655 targets in the MITLL SNP mixture panel were previously designed and vetted. Libraries were prepared using the AmpliSeq 2.0 library kit protocol according to the manufacturer recommendations, with the exception that 19 cycles were performed (no secondary amplification) and the library was eluted in 25uL low TE. Library quantitation was performed using the Ion Library Quantitation Kit (Thermo Fisher cat#4468802), according to the manufacturer recommendations. Template preparation and sequencing were performed using the Ion Chef and Ion Proton according to the manufacturer recommendations (Thermo Fisher Ion Chef and Proton cat#A27198 and Proton chips cat#A26771).

### MPS SNP Data Analysis

Ion Torrent BAM files were converted to FASTQ format using Samtools fastq(29). The GrigoraSNPs(24) program was used to call SNP alleles from multiplexed HTS FASTQ sequences. Mixture analysis was performed using the MIT Lincoln Laboratory IdPrism HTS DNA Forensics system. IdPrism uses the FastID(30) program to compare mixtures to reference samples. The IdPrism Plateau(18) method can also identify individual SNP profile signatures by mixture deconvolution. IdPrism uses the Fast P(RMNE)(31) program to automatically calculate the probability of a random person (man) not excluded P(RMNE)(16, 17). For saturated (i.e., > 70% of loci detected with minor alleles) SNP mixtures, IdPrism uses the TranslucentID(32) program to identify individuals in saturated mixtures that are not detected using standard mixture analysis techniques. The TranslucentID method is a novel approach to the identification of individuals in a saturated mixture that creates a derivative desaturated mixture by treating the SNPs with the lowest mAR values as MM alleles(32). This method was used on the equimolar ten person defined mixture (URK5V:IX-25).

EuroForMix(v1.11.4)(3, 4) was used to calculate continuous maximum likelihood ratios for each subject identified for mixtures with five or fewer contributions. The R program, Grigora_to_euroformix.R, was used to parse GrigoraSNPs output files into EuroForMix input format. EuroForMix settings were either tailored to the MITLL mixture panel or set at levels described in Bleka et al.(3) For each likelihood ratio the following hypotheses were compared:

H1: The person of interest and N-1 unknown individuals contributed to the mixture
H2: N unknown individuals contributed to the mixture

### Mean and Median Minor Allele Ratios Methods

The most common SNP allele in a population is referred to as the major (M) allele. The other SNP alleles are referred to as the minor (m) allele(s). The majority of SNP loci have predominantly two alleles. The minor allele ratio (mAR) at a locus is defined as the ratio of minor allele sequence counts divided by the total sequence counts. The minor allele ratio (mAR) for alleles in a SNP reference profile are approximately 0.0 for MM alleles (genotypes), a normal distribution centered on 0.5 for mM alleles, and at or near 1.0 for mm alleles. The loci with mM SNP alleles for an individual within a mixture reflect the relative concentration of the DNA contributed – i.e., the mAR of minor alleles contributed by only one individual will be represented in relative proportion to the contributor’s DNA in the sample compared to all other major alleles at each of these loci. When two or more minor alleles are present at a locus, the mAR approximate the additive combination of the contributors’ individual loci mAR values.

When reference profiles or Plateau(18) method profiles are available, the average or median of the unique mM alleles contributed to a mixture can be used to estimate the relative contributions of the individual’s DNA to the mixture. For a reference profile, the mAR for the mM alleles center in a normal distribution around 0.5 (with MM at 0 and mm at 1). The expected value for each DNA contributor should be directly proportional to their concentration in a mixture of two or more individuals. Figure 1 illustrates the mixture profile for a mixture with 5 contributors. The minor alleles for each contributor to this mixture are shown in Figure 2. SNP loci with minor alleles shared by two or more individuals are excluded from the calculations for known contributors with reference profiles. This is illustrated in Figure 3 (unique minor alleles) compared to Figure 2 (unique and shared minor alleles). The concentration of each contributor’s DNA is then estimated as 2 * *μ*, where *μ* is either the mean or the median mAR of the SNP loci where the given individual uniquely contributes a minor allele to the mixture (Figure 3 diamond marker for *μ*). For mixtures with unidentified contributors, unassigned mM alleles are leveraged to estimate the concentration for remaining unknowns in the mixture.

**FIG. 1.**
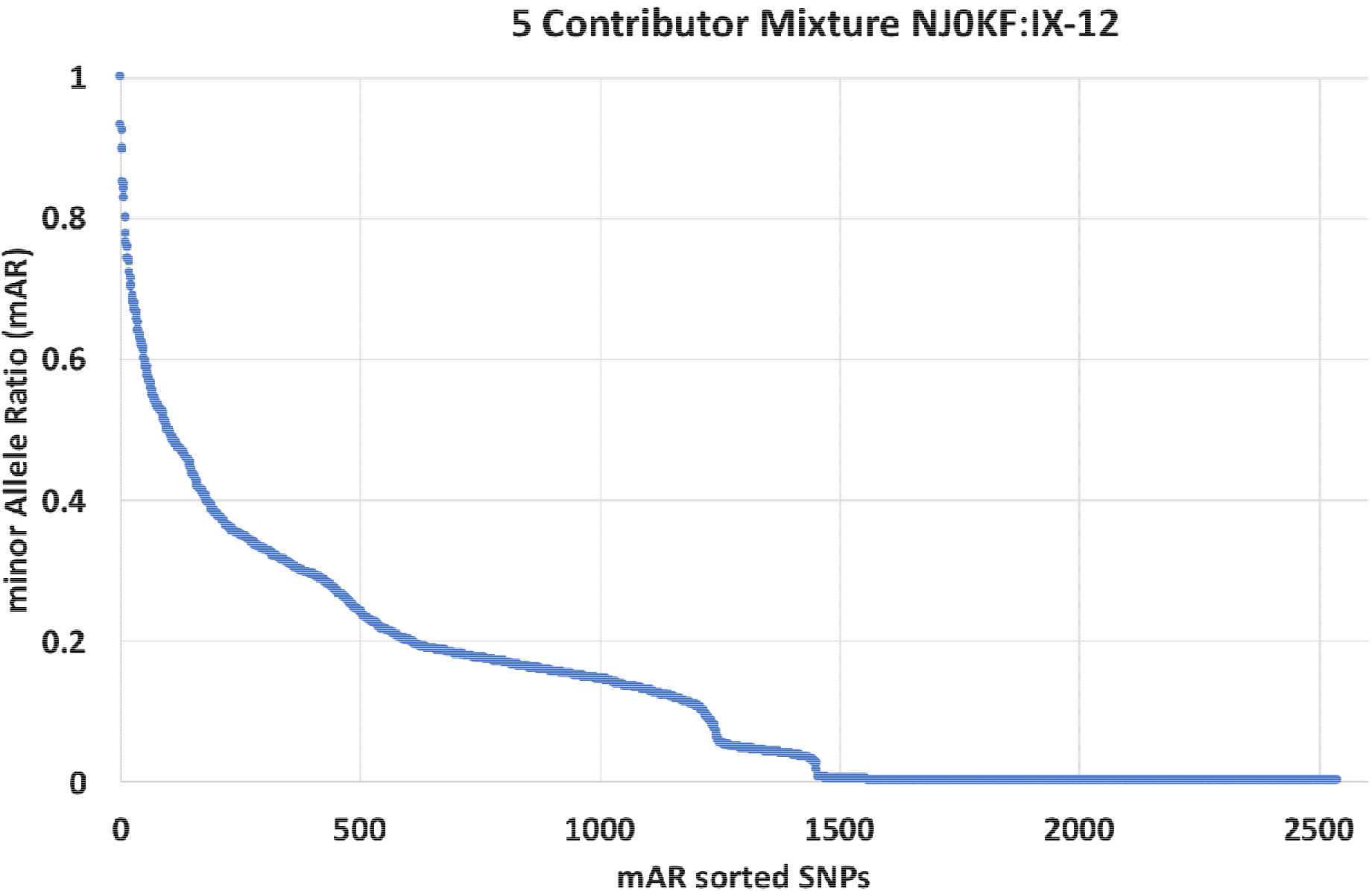
Example five-person mixture (NJ0KF:IX-12) illustrating mixture profile sorted by decreasing minor Allele Ratio (mAR).

**FIG. 2.**
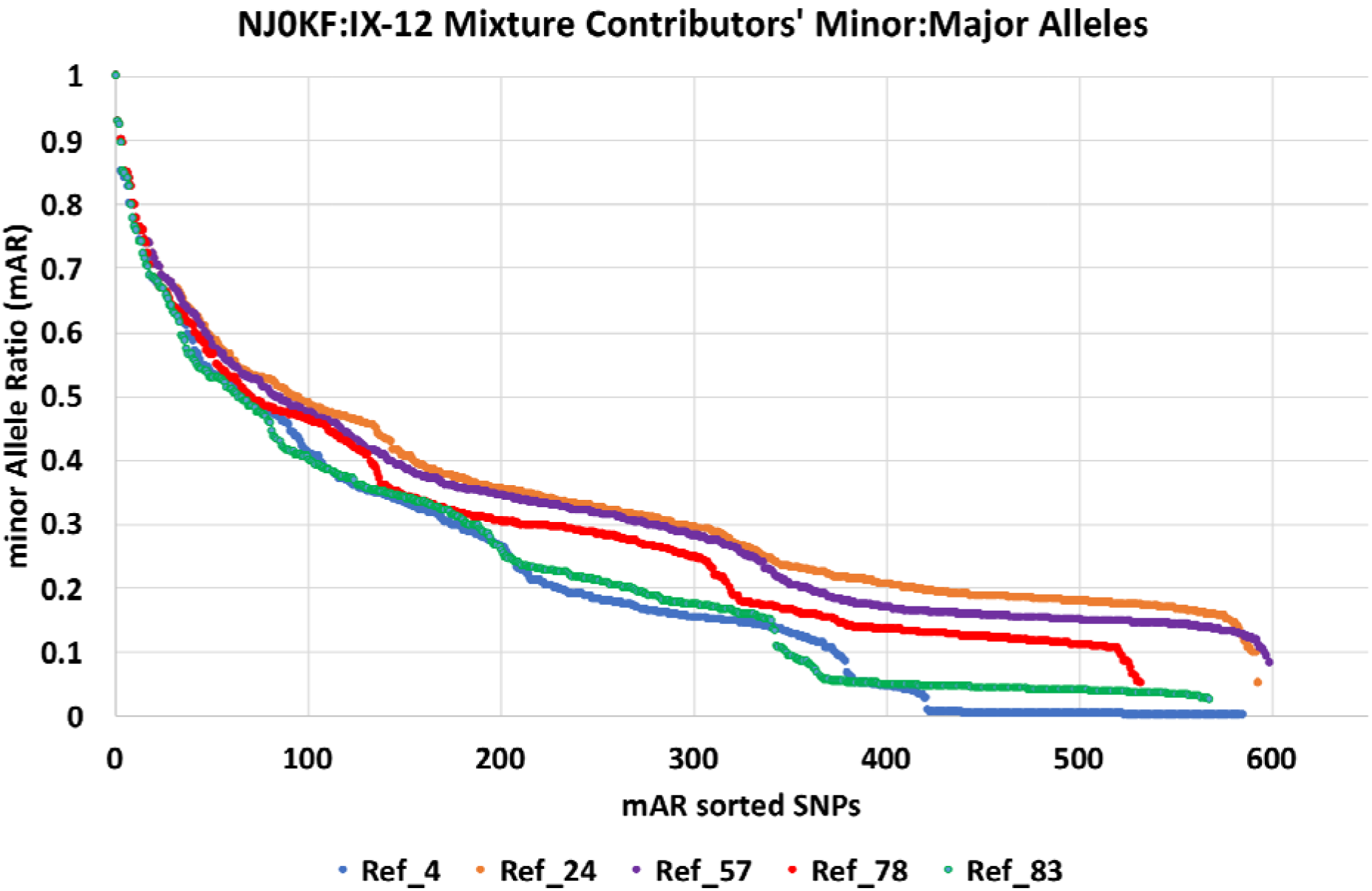
Mixture profile for all minor alleles of identified contributors to five-person mixture (NJ0KF:IX-12) sorted by decreasing mAR.

**FIG. 3.**
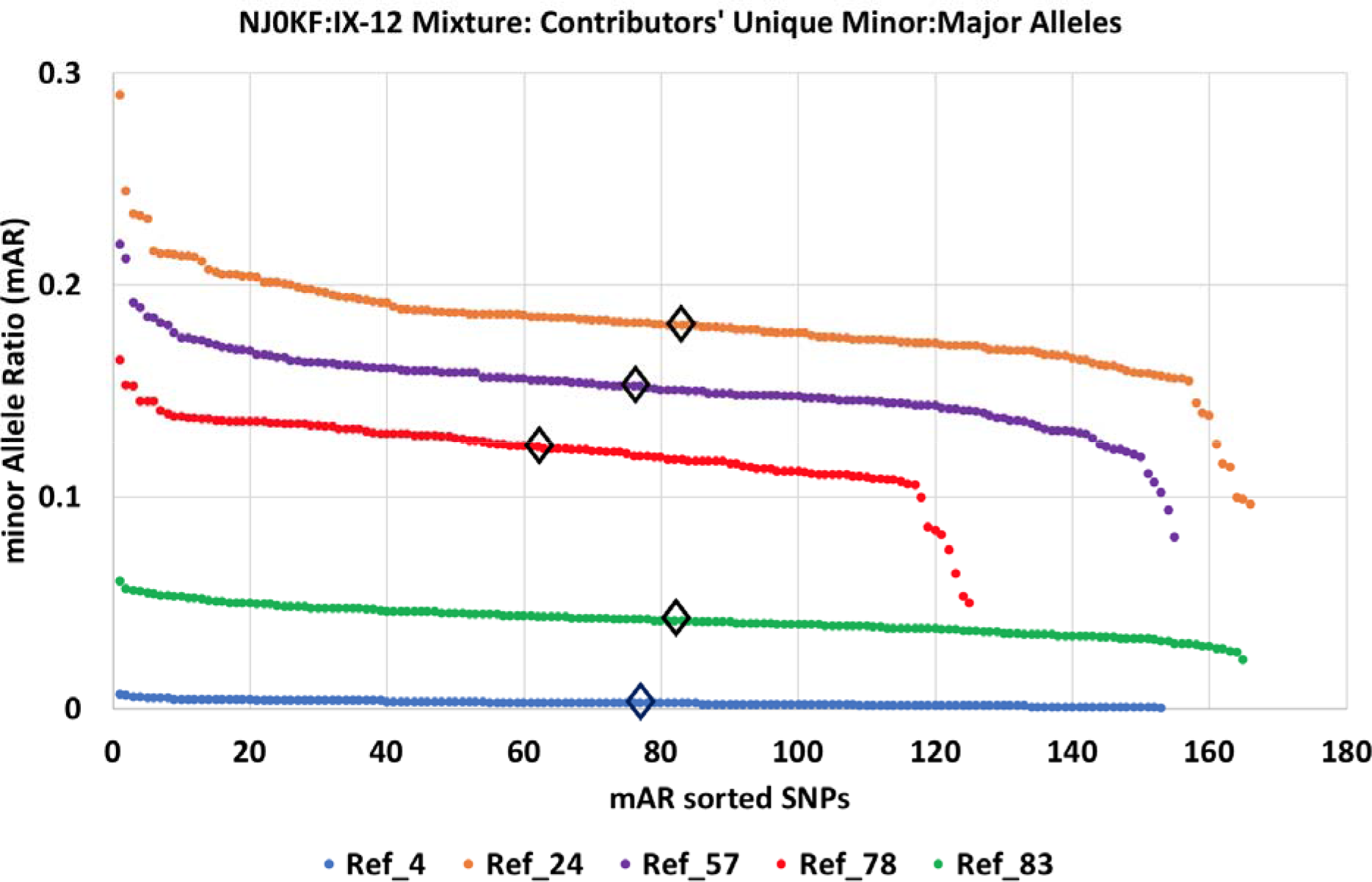
Estimating contributor concentrations from medium (diamond markers) or mean of minor alleles contributed uniquely by each contributor to five-person mixture (NJ0KF:IX-12).

### Slope Intercept Method

Unique mM SNPs for each individual in a mixture are determined by FastID(30) or the Plateau method(18). A linear regression (R language lm function) is performed with mAR as the dependent variable and each SNP as an independent variable. The slope intercept of each mAR regression is summed. Each individual slope intercept is divided by the sum of slope intercepts to determine individual DNA concentrations. Figure 4 illustrates the y-axis slope intercepts for a 5-person mixture on unique mM SNPs for each contributor.

**FIG. 4.**
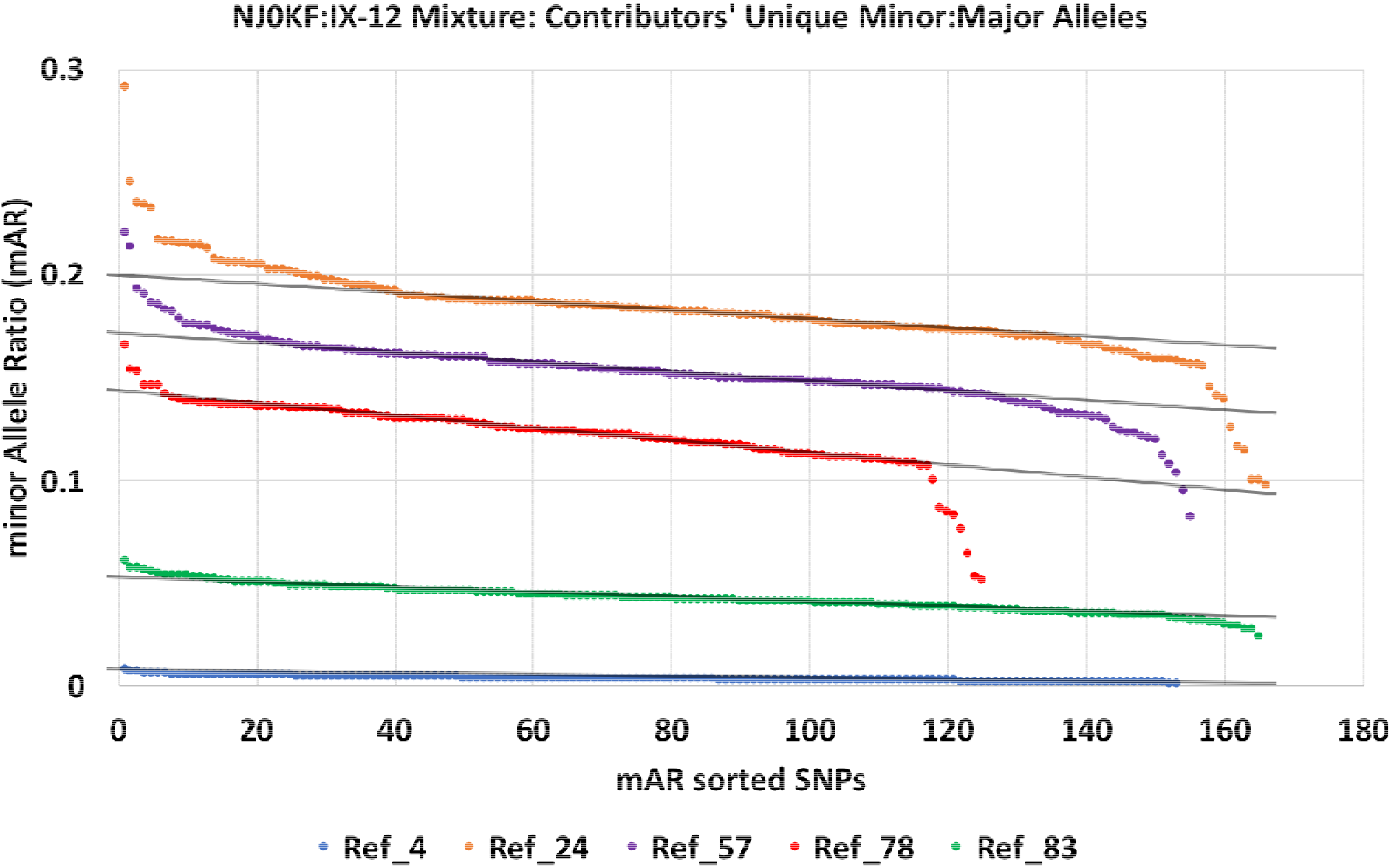
Y-axis slope intercepts based on contributor’s unique minor alleles for estimating mixture contributor concentration to five-person mixture (NJ0KF:IX-12).

## Results

### Defined Mixtures

Analysis results of defined mixtures of two to five individuals are illustrated in Table 1, where the estimated contributions from all three methods are compared to the intended contributions (the planned concentrations). The difference between the estimated concentration and the planned concentration is typically within 5%, with the exception of experiment UYLML:IX-31. For this four-person mixture, the methods consistently estimate more DNA from individual 93 and less DNA from individual 94 than was intended. Experimental error was identified after analysis of mixtures 5Z1MO:IX-01, IX-02, and IX-03. Table 1 reflects the revised planned concentration values for these three experiments.

**Table 1.**
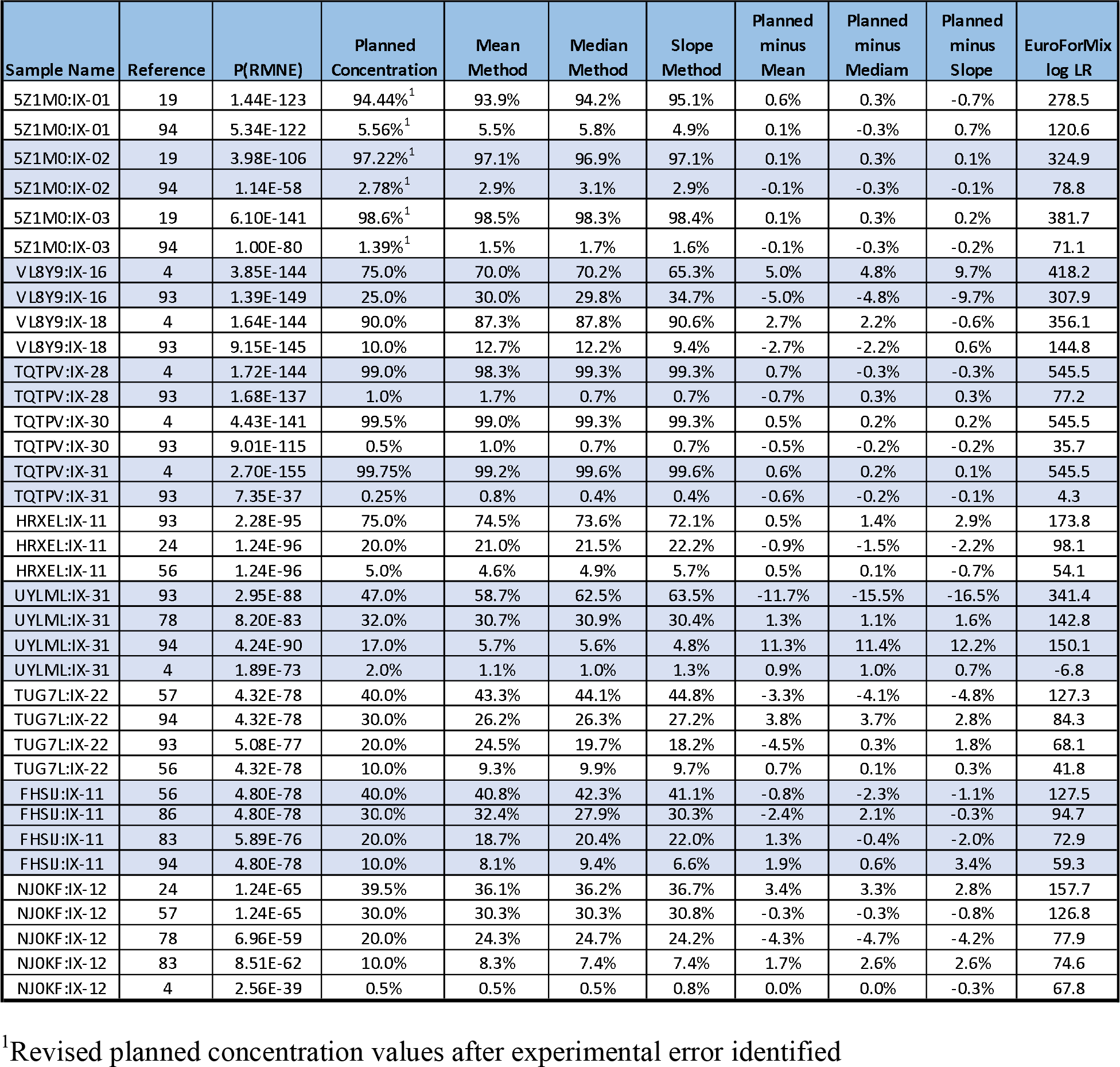
Estimation of contributions compared with intended percent contribution

### Example Saturated Equimolar Mixture

An equimolar mixture was created with DNA from ten individuals. This number of DNA contributors saturated the MITLL SNP mixture panel. By default, IdPrism does not perform standard mixture analysis on saturated mixtures. The MITLL TranslucentID method was applied to desaturate the mixture. FastID(30) identifies nine of the ten individuals in the desaturated mixture, see Table 2.

**Table 2.** Estimation of contributions for desaturated 10 individuals equimolar mixture (URK5V:IX-25), individual 41 was not detected (ND).

| Reference | P(RMNE) | Target<sup>?</sup><br>Concentration | Mean<sup>?</sup><br>Method | Median<sup>?</sup><br>Method | Slope<sup>?</sup><br>Method | Planned<sup>?</sup><br>minus<sup>?</sup><br>Mean | Planned<sup>?</sup><br>minus<sup>?</sup><br>Mediam | Planned<sup>?</sup><br>minus<sup>?</sup><br>Slope |
| --- | --- | --- | --- | --- | --- | --- | --- | --- |
| 4 | 1.30E-25 | 10% | 10.27% | 10.03% | 13.15% | -0.3% | 0.0% | -3.2% |
| 24 | 3.05E-28 | 10% | 10.31% | 10.56% | 7.99% | -0.3% | -0.6% | 2.0% |
| 41 | ND | 10% | - | - | - | - | - | - |
| 57 | 3.05E-28 | 10% | 9.37% | 9.34% | 9.46% | 0.6% | 0.7% | 0.5% |
| 68 | 3.05E-28 | 10% | 10.30% | 10.39% | 9.05% | -0.3% | -0.4% | 0.9% |
| 69 | 3.05E-28 | 10% | 10.01% | 10.19% | 12.96% | 0.0% | -0.2% | -3.0% |
| 76 | 3.05E-28 | 10% | 11.35% | 11.70% | 8.70% | -1.4% | -1.7% | 1.3% |
| 78 | 1.30E-24 | 10% | 9.48% | 9.39% | 13.53% | 0.5% | 0.6% | -3.5% |
| 83 | 1.30E-24 | 10% | 9.81% | 9.60% | 9.07% | 0.2% | 0.4% | 0.9% |
| 93 | 1.15E-27 | 10% | 12.60% | 12.13% | 12.57% | -2.6% | -2.1% | -2.6% |

### Example Visualizations

Figures 5 and 6 illustrate the mean method applied to a defined mixture and a mixture created from individuals touching an object. While there are no known concentrations for the touch mixture, the number of people and touch order was recorded. Mixture analysis with FastID(30) and reference profiles identifies five of the six individuals who touched this object (wood). Individual 83 was not detected in the mixture, but may be partially included in the signature marked “unknowns”. Table 3 estimates the relative contributions to the mixture by the individuals using the mean method.

**Table 3.**
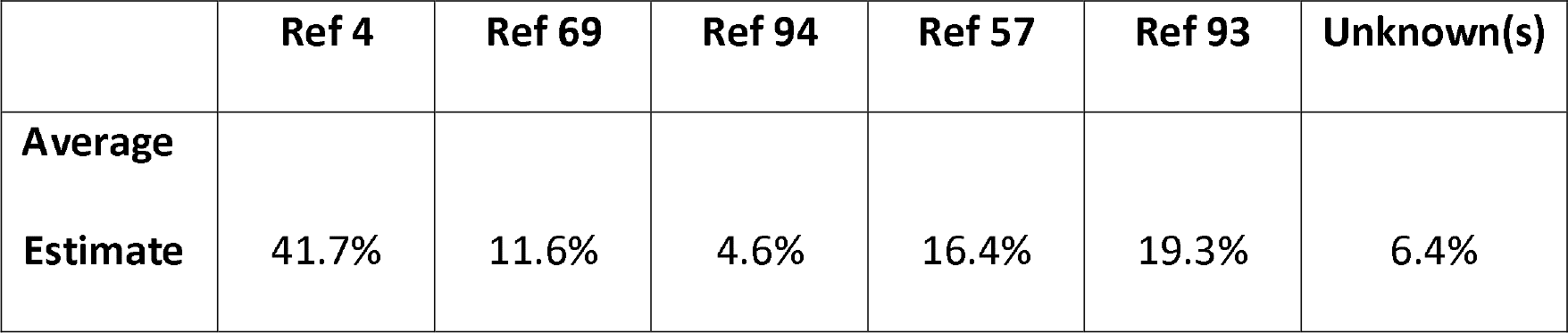
Estimated contributor concentrations for touch mixture mix160

**FIG. 5.**
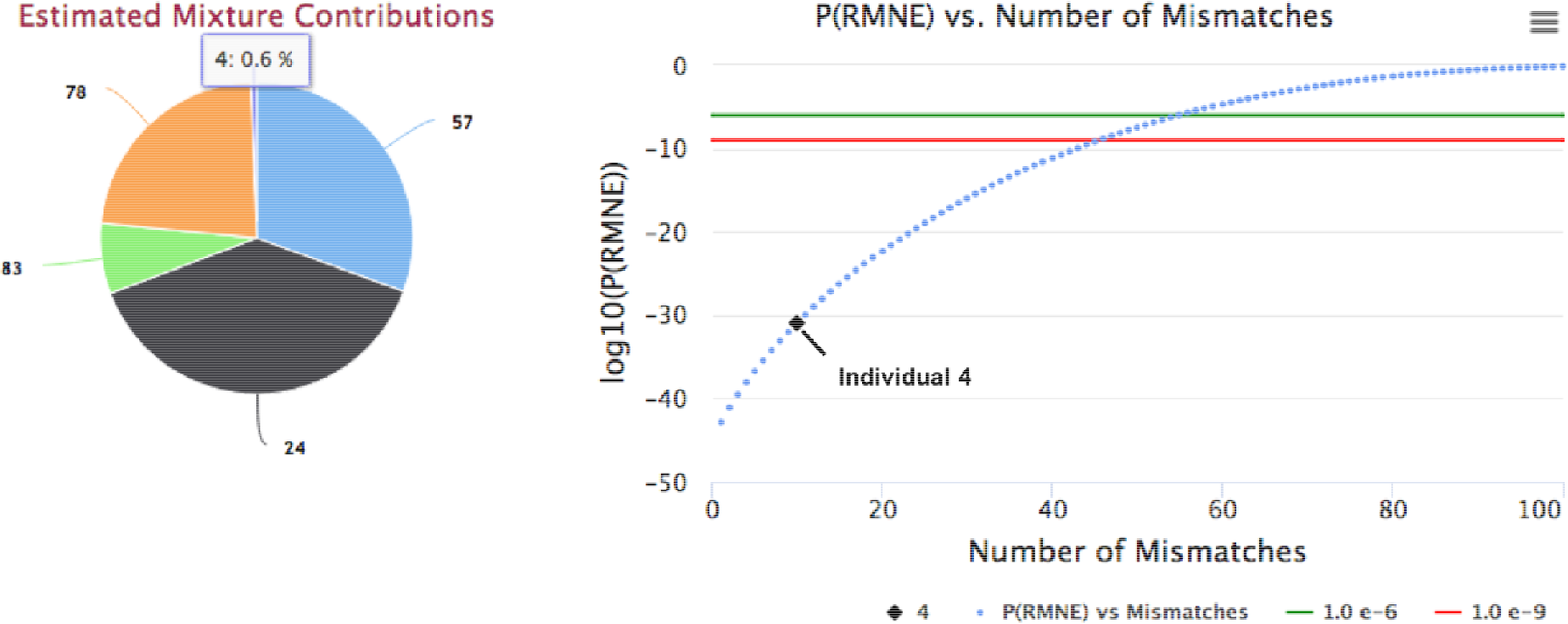
IdPrism screenshot of Five person mixture (NJ0KF:IX-12) with reference 4 selected in the Pie chart and displayed on the companion P(RMNE) (31) graph. Individual 4 has 17 dropped SNP minor alleles not detected above the analytical threshold(18); individuals 24, 57, 78, and 83 have no dropped minor alleles.

**FIG. 6.**
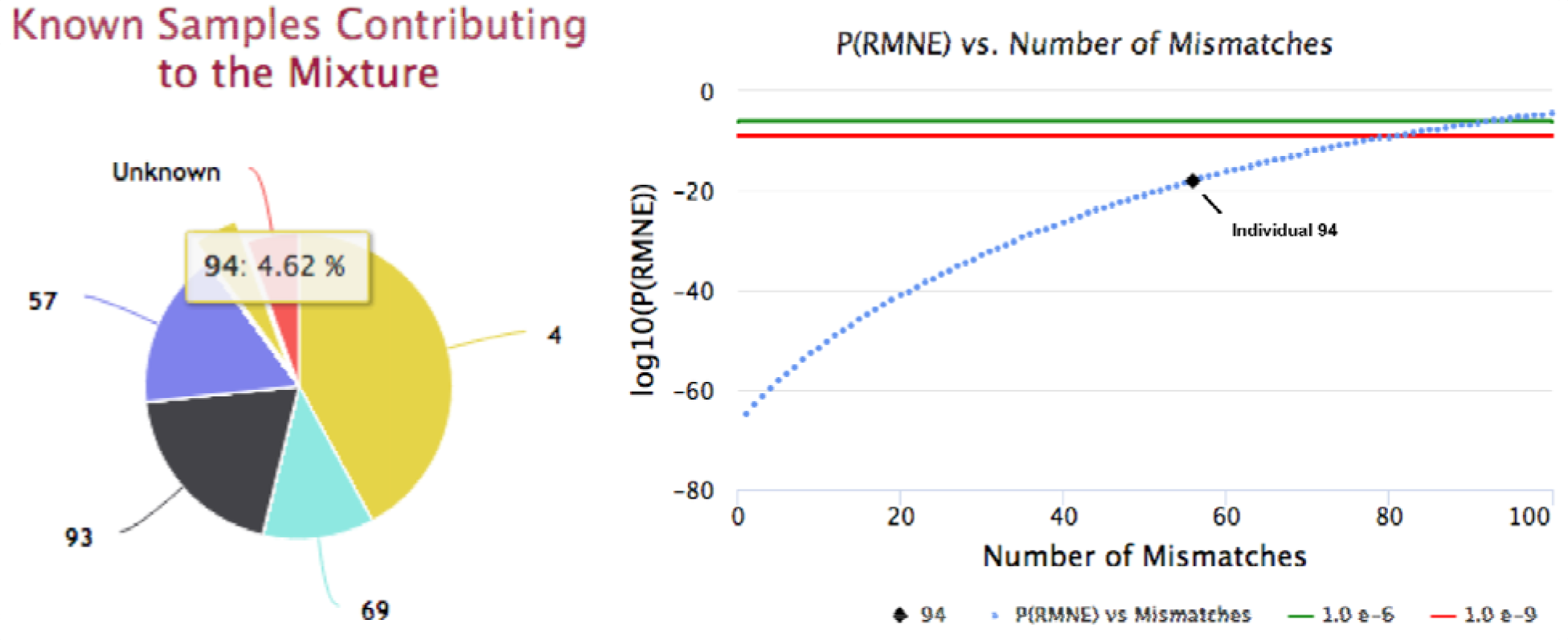
IdPrism screenshot of a touch mixture (09N7W:IX-29) with reference 94 selected (yellow slice) in the Pie chart and displayed on the companion P(RMNE) graph. This touch mixture had six known DNA contributors: 4, 57, 69, 83, 93, and 94 of which five contributors are detected with high confidence. Highlighted individual 94 has a concentration of 4.58% and a P(RMNE) of 1.05e-18, with 56 dropped mM loci not detected above the mixture analytical threshold.

## Discussion

Estimating the relative contributions of contributors in SNP mixtures is analogous to using STR peak heights or areas; with sharing of alleles by contributors a confounder for both data types. Focusing on the unique mM alleles contributed by each individual provides fairly accurate predictions with the variance from intended experimental planned concentrations in defined mixtures typically less than 5% for all three methods, see Table 1. These methods work for DNA contributor concentrations as low as 1:400 (TQTPV:IX-31 in Table 1) and mixtures with large numbers of contributors, (Table 2). The combined contributions for unknown individuals can also be estimated (Figure 6 and Table 3). Observed variation may reflect minor inaccuracy of source DNA concentration assessment, pipetting variability, or other mitigating factors. The IdPrism Pie chart visualization provide immediate insights into mixture contributors and unknowns. The inclusion of the mean method into MITLL IdPrism HTS Forensic System enabled the immediate identification of discrepancies in the intended concentrations of 5Z1MO:IX-01 of 1:200 at 1:18, 5Z1MO:IX-02 of 1:400 as closer to 1:34, and 5Z1MO:IX-03 of 1:800 as closer to 1:67. Detection of an experimental error enabled the correction of the planned concentrations of these experiments to 1:18, 1:36, and 1:72 as shown in Table 1. These updated concentrations are in close agreement with the values estimated by these methods.

These methods are also useful for DNA mixtures with no *a priori* known concentrations. For the touch example (09N7W:IX-29), individual 4 contributed more DNA to the object even though individual 93 was the last documented individual to touch the object. It has been previously established that the amount of DNA contributed by touch varies by individual(33). Figure 6 illustrates the estimated concentrations for the 5 detected individuals and the amount contributed by unknown individuals estimated at 6.3%.

## Conclusion

Three methods were introduced for estimating DNA contribution in mixtures characterized by HTS SNP panels. The values estimated by these methods are typically within 5% for defined DNA mixtures. These methods can also estimate the remaining amount of DNA contributed from unidentified individuals (unknowns).

## Acknowledgements

The authors acknowledge Sara Stankiewicz for useful conversations.

### DISTRIBUTION STATEMENT A.

Approved for public release. Distribution is unlimited.

This material is based upon work supported under Air Force Contract No. FA8702-15-D-0001. Any opinions, findings, conclusions or recommendations expressed in this material are those of the author(s) and do not necessarily reflect the views of the U.S. Air Force.

## References

1. Oldoni F, Castella V, Hall D. Shedding light on the relative DNA contribution of two persons handling the same object. Forensic Sci Int Genet. 2016;24:148–57.

2. Kanokwongnuwut P, Martin B, Kirkbride KP, Linacre A. Shedding light on shedders. Forensic Sci Int Genet. 2018;36:20–5.

3. Bleka Ø, Storvik G, Gill P. EuroForMix: An open source software based on a continuous model to evaluate STR DNA profiles from a mixture of contributors with artefacts. Forensic Sci Int Genet. 2016;21:35–44.

4. Bleka Ø, Eduardoff M, Santos C, Phillips C, Parson W, Gill P. Using EuroForMix to analyse complex SNP mixtures, up to six contributors. Forensic Sci Int Genet. 2017;6(Genet Suppl):e277–e9.

5. Bill M, Gill P, Curran J, Clayton T, Pinchin R, Healy M, et al. PENDULUM—a guideline-based approach to the interpretation of STR mixtures. Forensic Sci Int. 2005;148(2):181–9.

6. Gill P, Sparkes R, Pinchin R, Clayton T, Whitaker J, Buckleton J. Interpreting simple STR mixtures using allele peak areas. Forensic Sci Int. 1998;91(1):41–53.

7. Cowell RG, Graversen T, Lauritzen SL, Mortera J. Analysis of forensic DNA mixtures with artefacts. J R Stat Soc Ser C Appl Stat. 2015;64(1):1–48.

8. http://www.armedxpert.com (accessed 2018).

9. Cowell RG, Graversen T, Lauritzen SL, Mortera J. Analysis of forensic DNA mixtures with artefacts. J R Stat Soc Ser C Appl Stat. 2015.

10. Brenner CH. The DNA VIEW Mixture Solution. ISFG2015.

11. https://www.qualitype.de/en/solutions/products/evaluation-software/genoproof-mixture/ (accessed 2018).

12. Inman K, Rudin N, Cheng K, Robinson C, Kirschner A, Inman-Semerau L, et al. Lab Retriever: a software tool for calculating likelihood ratios incorporating a probability of drop-out for forensic DNA profiles. BMC Bioinformatics 2015;16(1):298.

13. Haned H, Slooten K, Gill P. Exploratory data analysis for the interpretation of low template DNA mixtures. Forensic Sci Int Genet. 2012;6(6):762–74.

14. Taylor D, Bright J-A, Buckleton J. The interpretation of single source and mixed DNA profiles. Forensic Sci Int Genet. 2013;7(5):516–28.

15. Perlin MW, Legler MM, Spencer CE, Smith JL, Allan WP, Belrose JL, et al. Validating TrueAllele^®^ DNA Mixture Interpretation. J Forensic Sci. 2011;56(6):1430–47.

16. Voskoboinik L, Darvasi A. Forensic identification of an individual in complex DNA mixtures. Forensic Science International: Genetics. 2011;5(5):428–35.

17. Isaacson J, Schwoebel E, Shcherbina A, Ricke D, Harper J, Petrovick M, et al. Robust detection of individual forensic profiles in DNA mixtures. Forensic Sci Int Genet. 2015;14:31–7.

18. Ricke DO, Isaacson J, Watkins J, Fremont-Smith P, Boettcher T, Petrovick M, et al. The Plateau Method for Forensic DNA SNP Mixture Deconvolution. bioRxiv. 2017.

19. Shcherbina A, Ricke DO, Schwoebel E, Boettcher T, Zook C, Bobrow J, et al. KinLinks: Software Toolkit for Kinship Analysis and Pedigree Generation from HTS Datasets. 2016 IEEE International Symposium on Technologies for Homeland Security (HST). 2016 May 10–11.

20. Helfer BS, Fremont-Smith P, Ricke DO. The Genetic Chain Rule for Probabilistic Kinship Estimation. bioRxiv. 2017.

21. Pakstis AJ, Speed WC, Fang R, Hyland FCL, Furtado MR, Kidd JR, et al. SNPs for a universal individual identification panel. Human Genet. 2010;127(3):315–24.

22. Nievergelt CM, Maihofer AX, Shekhtman T, Libiger O, Wang X, Kidd KK, et al. Inference of human continental origin and admixture proportions using a highly discriminative ancestry informative 41-SNP panel. Investigative Genet. 2013 2013;4:13.

23. Branicki W, Liu F, van Duijn K, Draus-Barini J, Pośpiech E, Walsh S, et al. Model-based prediction of human hair color using DNA variants. Hum Genet. 2011 2011;129(4):443–54.

24. Ricke DO, Shcherbina A, Michaleas A, Fremont-Smith P. GrigoraSNPs: Optimized HTS DNA Forensic SNP Analysis. J Forensic Sci. 2018.

25. http://www.illumina.com/areas-of-interest/forensic-genomics/forensic-analysis-methods/snp-str-analysis.html (accessed 2018).

26. https://www.battelle.org/government-offerings/homeland-security-public-safety/security-law-enforcement/forensic-genomics/exactid/technical-specifications (accessed 2018).

27. Woerner AE, King JL, Budowle B. Fast STR allele identification with STRait Razor 3.0. Forensic Sci Int Genet. 2017;30(Suppl C):18–23.

28. Ricke DO, Harper J, Helfer BS, Isaacson J, Michaleas AM, Petrovick MS, et al., inventors; Massachusetts Institute of Technology, assignee. Systems and Methods for Genetic Identification and Analysis. International Patent Application PCT/US2018/041081 (2018, July 6).

29. Li H, Handsaker B, Wysoker A, Fennell T, Ruan J, Homer N, et al. The Sequence Alignment/Map format and SAMtools. Bioinformatics. 2009;25(16):2078–9.

30. Ricke DO. FastID: Extremely Fast Forensic DNA Comparisons. IEEE Explore. 2017. 31.

31. Ricke D, Schwartz S. Fast P(RMNE): Fast forensic DNA probability of random man not excluded calculation [version 1; referees: awaiting peer review]. F1000Research. 2017;6:2154.

32. Ricke DO, Watkins J, Fremont-Smith P, Boettcher T, Schwoebel E. TranslucentID: Detecting Individuals with High Confidence in Saturated DNA SNP Mixtures. bioRxiv. 2018.

33. Lowe A, Murray C, Whitaker J, Tully G, Gill P. The propensity of individuals to deposit DNA and secondary transfer of low level DNA from individuals to inert surfaces. Forensic Sci Int. 2002;129(1):25–34.

